# Ablation and Seed-driven Restoration of an Alpha-Satellite Devoid Human Centromere Reveals Size Homeostasis Mechanisms

**DOI:** 10.64898/2026.08.14.744943

**Authors:** Miao Lin, Shan Hua, Cody J. Naccarato, Zhao Zhang, Zhijie Liu, Fen Guo, Vanessa Horner, M. Anwar Iqbal, Archibald Perkins, Patrick J. Murphy, Bin Zhang

## Abstract

Human centromeres are epigenetically defined chromatin domains marked by nucleosomes containing the histone H3 variant CENP-A, which recruit the constitutive centromere-associated network (CCAN) to assemble functional kinetochores. Maintaining centromere function, including chromatin domain size and the ability to assemble the kinetochore, is essential for proper mitotic division across eukaryotes. In humans however, mechanistic studies of centromere establishment, maintenance, and size regulation have been hindered by the highly repetitive nature of canonical alphoid centromeres. Here, we develop a genetically tractable human neocentromere system which is devoid of repetitive DNA sequences. Using targeted genetic manipulation of a monoallelic naturally occurring neocentromere, we show that partial loss of centromeric chromatin triggers restoration of the residual CENP-A domain through seed-driven, sequence-independent expansion into adjacent naïve chromatin. In contrast, creation of new domain boundaries without loss of centromeric chromatin mass results only in local boundary remodeling, without substantial domain expansion. These findings indicate that centromere formation proceeds through two mechanistically distinct steps, beginning with acquisition of a CENP-A seed and followed by regulated domain expansion to generate a mature centromere. More broadly, our results support a model in which centromeres transition between a maintenance state that preserves domain size and a restoration state that rebuilds centromeric chromatin following perturbation. Together, this study establishes a genetically tractable platform for dissecting the mechanisms governing human centromere formation, chromatin domain dynamics, and size homeostasis.

## INTRODUCTION

Centromeres are essential chromatin structures that specify the kinetochore assembly location on each chromosome. Properly functioning centromeres recruit at least 16 centromere proteins (CENPs), which comprise the <u>c</u>onstitutive <u>c</u>entromere-<u>a</u>ssociated <u>n</u>etwork (CCAN), also known as the inner kinetochore. The CCAN in turn recruits the outer kinetochore of the KMN network (made up of KNL1, MIS12, and NDC80), to which the microtubule spindle attaches during mitosis and meiosis^1–3^. This step-wise recruitment is critical for accurate chromosome segregation during cell division. At the cellular level, dysregulation of centromeres causes aneuploidy and/or chromosomal instability, leading to alteration of cell fitness or cell death^4^. At the organismal level, germline chromosome non-disjunction that result from compromised centromere activity can have far-reaching consequences for development, affecting the viability of pregnancies as well as causing developmental disorders such as Down syndrome. Moreover, somatic alterations impacting centromere function can promote both tumor initiation and progression^4–6^.

In human cells, canonical centromeric chromatin is typically associated with repetitive α-satellite DNA (alphoid)^7,8^. However, centromere identity is not determined by DNA sequence alone, and instead, epigenetic chromatin features establish centromere identity, most notably nucleosomes containing the histone H3 variant, CENtromere Protein A (CENP-A), which nucleate kinetochore assembly through interacting with CENP-C and CENP-N^1,2,9^. Two cytogenetic phenomena strongly indicate this sequence-independent centromere-specifying mechanism. First, centromeres on the alpha satellite repeats can lose CENP-A nucleosomes and become inactivated without changes to the underlying DNA sequences^10,11^. Second, fully functional centromeres can spontaneously arise at new genomic regions devoid of alpha satellites through mechanisms yet to be defined^5,10,12,13^.

Centromeres on non-repetitive sequences, neocentromeres, are known to stabilize disease-causing supernumerary marker chromosomes (SMCs), which include somatic giant ring chromosomes in subtypes of lipomatous tumors^14^, and germline inverted duplications of terminal regions of a chromosome, causing constitutional developmental disorders^15–17^. In particular, inv dup(15q), resulting in tetrasomy 15q25.2→qter, is among the most common neocentric marker chromosomes identified so far, causing a consistent phenotype including severe developmental delay, craniosynostosis, and overgrowth^15,18^. Although described as clinical cases in the literature, such marker chromosomes and the associated neocentromeres have not been extensively examined at the molecular or mechanistic level.

The molecular mechanisms underlying the formation, inactivation, and dysregulation of this epigenetic entity, functional CENP-A chromatin domain, remain as a main focus of biomedical research. Spontaneous formation of CENP-A chromatin domains within the genome do occur occasionally, but this is generally suppressed before forming functional neocentromeres, in order to protect genome integrity and prevent the catastrophic outcomes of chromosome loss and genome instability, resulting from possessing more than one centromere per chromosome^19^. For example, CENP-A nucleosomes ectopically loaded in the genome can be efficiently removed during DNA replication^19,20^. Such suppressive mechanisms are likely to negatively regulate random CENP-A chromatin domain establishment at ectopic sites and promote inactivation of one of the two centromeres on dicentrics.

The relative position of centromeres on each chromosome is remarkably stable when centromeres are epigenetically inherited during mitosis and meiosis. In this regard, CENP-A nucleosomes are efficiently recycled and inserted back into centromeric chromatin domain during DNA synthesis^21,22^. Deposition of new CENP-A nucleosomes onto diluted chromatin domain occurs in G1 and is regulated by the nucleosome loading chaperone Holliday Junction Recognition Protein (HJURP), the reloading licensing complex MIS18, and the reader of preexisting centromeric chromatin CCAN^23–27^. This cell-cycle-regulated inheritance of CENP-A chromatin follows a conservative template-guided replenishment mechanism, in which each pre-existing CENP-A nucleosome licenses the incorporation of one new CENP-A nucleosome to maintain domain size through cell cycles ^1,21,22,24,25,28^. Consistent with this, human centromeres appear to contain approximately 300-400 CENP-A molecules, with modest differences across cell types and little inter-centromere variability^29,30^. Together, both the position and size of human centromeres, as measured by the extent of the CENP-A chromatin domain, are epigenetically determined and relatively stable, even when the underlying DNA content varies^29–31^

The mechanisms that prevent centromere drift and regulate centromere size remain poorly understood^32^. Progress has been limited by the lack of systems to study the epigenetic establishment of centromeric chromatin and its expansion during formation or in response to damage. A major barrier to addressing these questions has been the repetitive nature of canonical centromeres^7,8^, which prevents precise genetic manipulation. Moreover, analysis of existing intact centromeres offers limited access to expansion dynamics of centromeric chromatin domain as its boundaries are restricted and its size are maintained through the conservative template-guided replenishment mechanism. Additionally, the organization, inheritance, and maintenance of centromeric chromatin are governed by rapidly evolving cis– and trans-acting components that exhibit substantial sequence divergence across species^7,33^, limiting the direct translation of findings from model systems to human cells and stressing the importance of developing new cell genetic systems to study human centromeric chromatin dynamics.

We have overcome these limitations by developing a genetically tractable human neocentromere system located on non-repetitive DNA, enabling direct and precise manipulation of centromeric chromatin in human cells. In this system, re-establishment of functional centromeric chromatin from defined chromatin seeds recapitulates a dynamic process that occurs sporadically *in vivo* during chromatin damage, *de novo* centromere formation, and subsequent maturation. By enabling inducible and synchronized seed-driven chromatin expansion, this platform allows direct interrogation of mechanisms governing centromere formation and size control. Notably, a transiently induced Active Domain Boundary (ADB), distinct from the restrictive boundary at intact centromeres, acts as a leading edge for new CENP-A deposition, providing a sensitive and tractable system to study domain expansion. This system delivers a useful framework to define key features of the local genetic and epigenetic landscape, as well as the essential protein factors, that govern centromere maintenance and chromatin domain boundary formation.

## RESULTS

### Clinical characterization of non-alphoid marker chromosomes reveals naturally occurring human neocentromeres

We first sought to identify naturally occurring alphoid-lacking human chromosomes which possessed neocentromeres, as these allow us to investigate the genetic and epigenetic mechanisms governing human centromere establishment and maintenance. We therefore searched for supernumerary marker chromosomes (SMCs) lacking canonical repetitive centromeric DNA in our clinical laboratories. We identified and analyzed a constitutional inv dup(15q) marker chromosome in an individual with multiple congenital anomalies. Chromosome microarray analysis demonstrated pathogenic tetrasomy of distal 15q, with the breakpoint mapping to a region enriched in segmental duplications (low-copy repeats) at 15q24.1– q24.3, a genomic architecture frequently associated with non-allelic homologous recombination (Fig. 1A). Interphase and metaphase fluorescence *in situ* hybridization (FISH) confirmed the duplicated genomic content and established the inverted duplication structure of the marker chromosome that lacks CEP15-specific alphoid sequences (Fig. 1B,C). We also identified and analyzed a somatic non-alphoid marker chromosome, inv dup(11q), causing hexasomy 11q in acute myeloid leukemia (Fig. 1D, E). Notably, metaphase FISH using the chromosome 11 alpha-satellite probe D11Z1 failed to detect alphoid sequences on the inv dup(11q) marker, despite robust hybridization to the normal chromosome 11 centromeres (Fig. 1F). Together, these findings demonstrate that both constitutional and somatically acquired marker chromosomes can exist in the absence of canonical alpha-satellite centromeres, indicating that they are stabilized by neocentromeres formed on non-repetitive DNA sequences. The rarity of such non-alphoid SMCs and their dependence on neocentromeres prompted us to develop an experimentally tractable model system for investigating the mechanisms that govern centromeric chromatin establishment, expansion, and domain size control.

**Figure 1:**
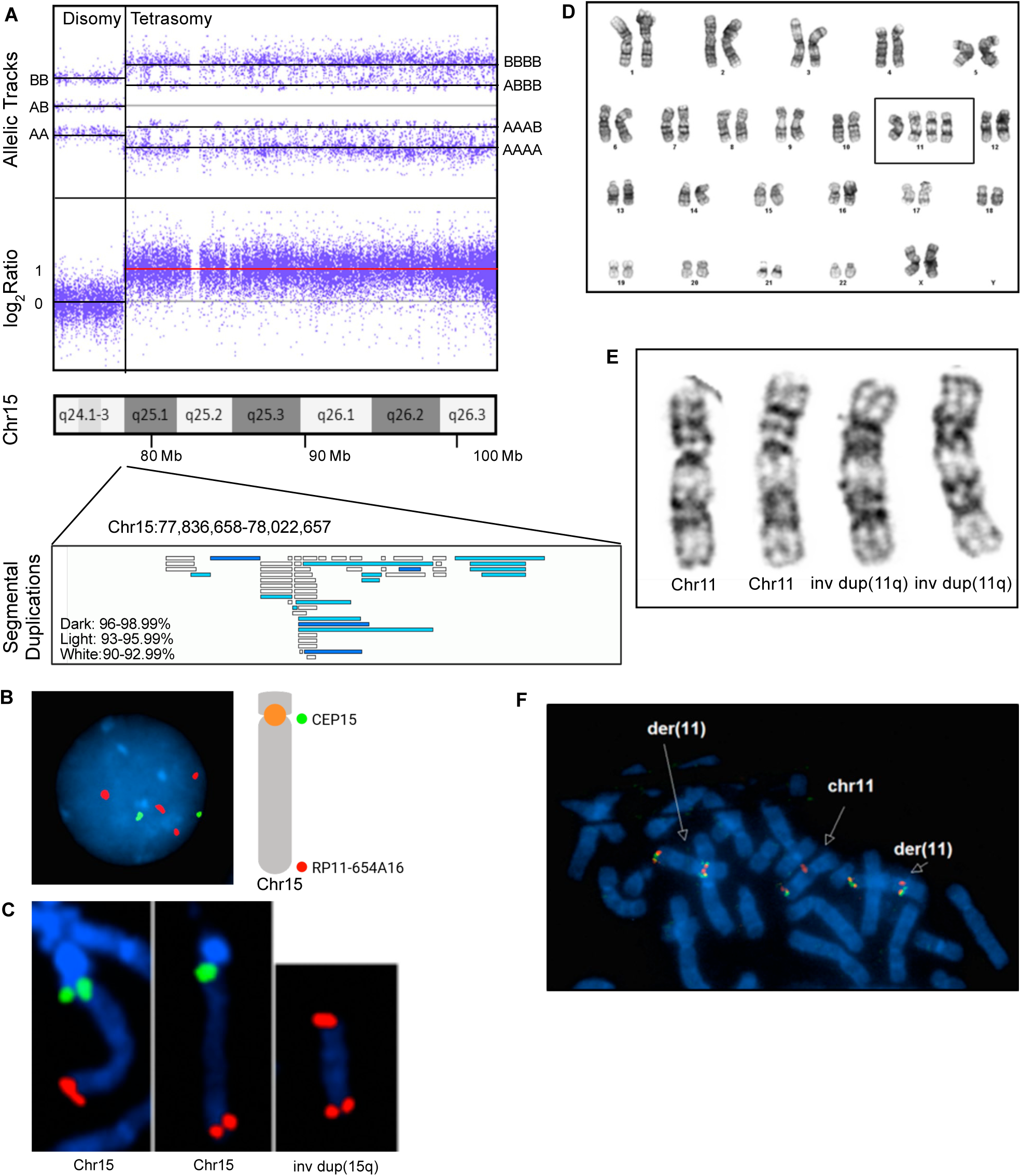
Cytogenetic characterization of marker chromosomes, inv dup(15q) (germline) and inv dup(11q) (somatic) from human patients, which lack alpha satellites and are likely stabilized by neocentromeres. **A)** Tetrasomy 15q distal to band 15q24.1-3 is detected in an individual affected by multiple congenital anomalies by chromosome microarray analysis (affymetrix). The breakpoint is mapped to a region comprised of low copy repeats, which likely contributed to the formation of a non-alphoid marker chromosome through non-allelic homologous recombination. **B)** Tetrasomy 15q is confirmed by interphase FISH analysis with the CEP15 (green) and RP11-654A16 (orange) probes. **C)** An inv dup(15q) marker chromosome results in tetrasomy 15q, confirmed by metaphase FISH. **D and E)** Two copies of an inv dup(11q) chromosome are detected in a bone marrow sample of acute myeloid leukemia (AML) using G-banded karyotyping analysis, (48,XX,inv dup(11q)x2). **F)** The inv dup(11q) does not harbor the alpha satellites of chromosome 11, evidenced with metaphase FISH using the D11Z1 (orange) and KMT2A (dual color dual fusion) probes.

### Development of a genetically tractable human neocentromere model

Rather than proceeding with these patient-derived samples, we next sought to establish a genetically tractable cellular models harboring naturally occurring neocentromeres. To this end, we searched the cell line database curated by the NIGMS Human Genetic Cell Repository at the Coriell Institute for Medical Research (Camden, NJ)^34^ using the terms: “i(15)”, “der(15)dup(15”, and “inv(15)”, and obtained GM09026 (a fibroblast cell line) and GM19715 (a lymphoblastoid cell line). These two lines were derived from individuals exhibiting overlapping congenital anomalies and carrying independently arising inv dup(15q) marker chromosomes, suggesting distinct neocentromere formation events.

We then characterized the GM09026 cell line genetically and epigenetically. G-banded chromosome analysis confirmed the presence of the marker chromosome in addition to two normal copies of chromosome 15 (Fig. 2A,B). Chromosome microarray analysis demonstrated tetrasomy of the distal 15q region and precisely mapped the breakpoint to 15q25.2, proximal to a region enriched for segmental duplications (Fig. 2C), the structure hypothesized to mediate the formation of inverted duplication chromosomes^15^. Combined immunofluorescence (IF) for CENP-A and fluorescence in situ hybridization (FISH) using a 15qter probe recognizing the subtelomeric region of chromosome 15 demonstrated that the marker chromosome harbors a functional centromere despite lacking the canonical chromosome 15 alphoid centromere (Fig. 2D). To define the location of the neocentromere, we performed CENP-A CUT&RUN sequencing and identified a prominent CENP-A-enriched domain at chromosome band 15q26.2, distinct from the endogenous centromere of chromosome 15 (Fig. 2E). Higher-resolution analysis revealed a major CENP-A-enriched domain spanning approximately 80 kb (NC1, hg38:chr15:94,666,054-94,746,161), flanked by two smaller satellite peaks located ∼25 kb from the main domain on either side (Fig. 2F). We refer to this non-repetitive centromeric chromatin domain as the CENP-A epi-allele (CA_epi), whereas other copies of the same 15q26.2 genomic region that lack CENP-A occupancy are designated non-centromeric alleles. Importantly, metaphase IF/FISH analysis demonstrated precise co-localization of CENP-A protein with the underlying 15q26.2 genomic sequence, confirming that the neocentromere is assembled on unique DNA rather than repetitive alphoid sequences (Fig. 2G). Together, these results identify a stable non-alphoid neocentromere located at 15q26.2 in the GM09026 line and establish a unique human genetic system in which centromeric chromatin can be directly manipulated at nucleotide resolution.

**Figure 2:**
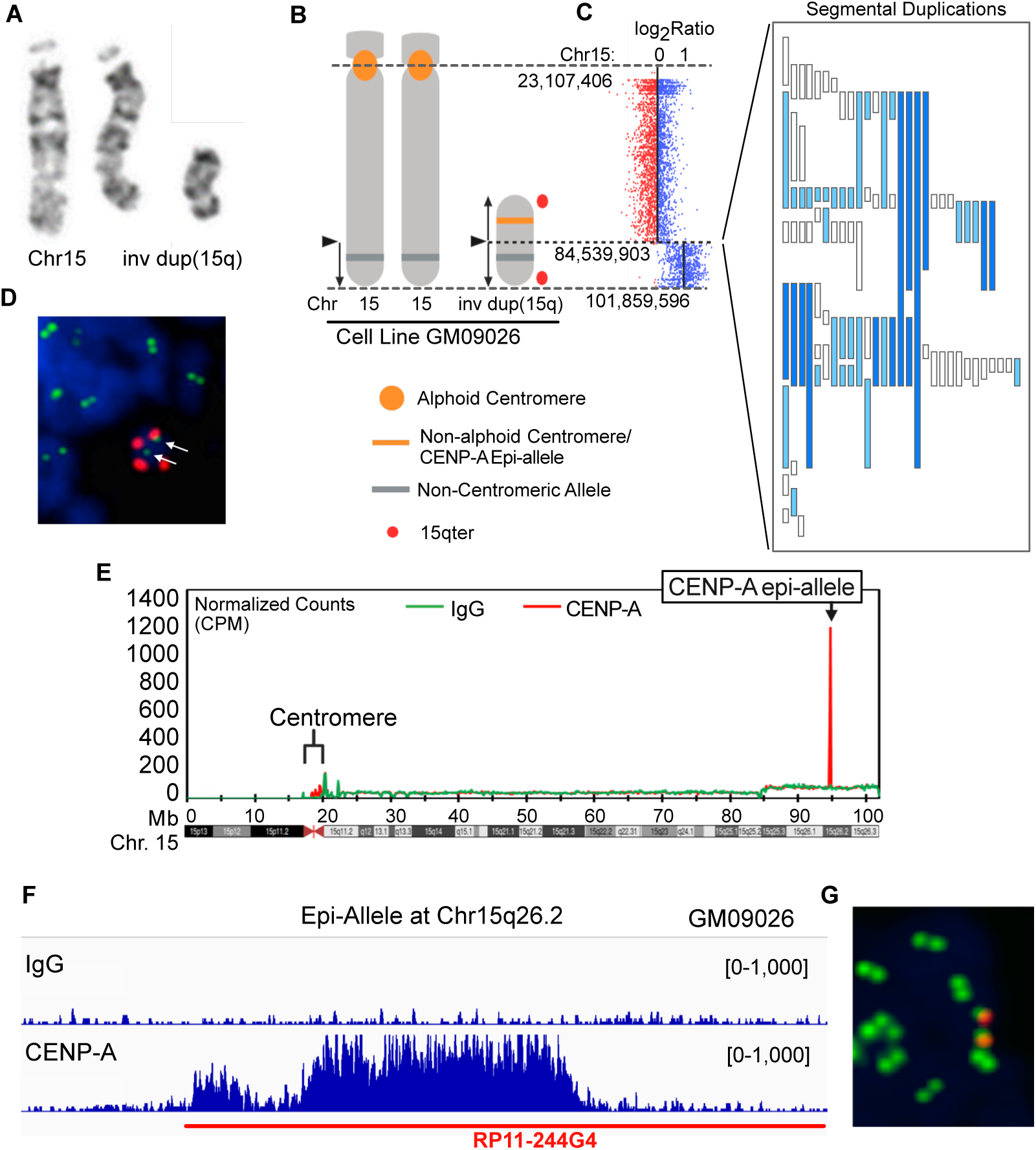
Genetic and epigenetic characterization of neocentromeres in human primary cell lines harboring non-alphoid markers. **A)** Cell line G026 derived from a human patient harbors an inv dup(15q), demonstrated by G-banded karyotyping analysis. **B)** Diagram of chromosome 15 and inv dup(15q). **C)** Chromosome microarray analysis (Agilent) defines the genomic content of tetrasomy 15q and genomic characteristics, a segmental duplication cluster distal to the breakpoint at 15q25.2. **D)** Metaphase IF/FISH analysis showed the inv dup(15q) harbors a non-alphoid centromere (CENP-A IF in green). **E)** CUT&RUN analysis with CENP-A antibody (red) and control IgG (green) defines the location of the neocentromere, referred to as a CENP-A epi-allele. It is noted that the less impressive enrichment of CENP-A at the canonical alphoid centromere on chromosome 15 is attributed to multiple mappers to alpha satellite repeats. **F)** A high-resolution view of the 15q26.2 CENP-A epi-allele. The location of RP11-244G4 is indicated. **G)** IF/FISH demonstrates co-localization of the CENP-A epi-allele (green) and the underlying non-alphoid DNA sequence (RP11-244G4 in red).

### Neocentromere landing regions are depleted of active chromatin marks and exhibit context-dependent enrichment of H3K9me3

Using CUT&RUN^35^ on GM19715, we defined a neocentromeric CENP-A chromatin domain spanning a 55 Kb region (NC2, hg38:chr15:93,681,637-93,736,985), about 1 Mb distal to the location of the neocentromere in GM09026 (NC1). These two neocentromere landing regions (NC1 and NC2) are noted for their significantly higher AT content relative to a surrounding 3 Mb region (59.3% and 60.7% vs. 50%). To investigate whether neocentromere formation is associated with distinctive chromatin environments, we analyzed publicly available epigenomic datasets from human fibroblast (IMR90), embryonic stem cell (H1), and leukemia (K562) cell lines, comparing these two independently identified neocentromere landing regions (NC1 from GM09026 and NC2 from GM19715), with canonical centromeres, gene promoters, or randomly selected genomic regions. As expected, the active chromatin marks H3K4me3 and H3K27ac were strongly enriched at gene promoters but were significantly depleted at both canonical centromeres and neocentromere landing regions (Fig. 3). In contrast, the facultative heterochromatin mark H3K27me3 showed no consistent enrichment or depletion among centromeres, neocentromeres, promoters, and random genomic regions across the cell lines examined (Fig. 3). The constitutive heterochromatin mark H3K9me3 displayed a more complex pattern. In IMR90 fibroblasts, both canonical centromeres and neocentromere landing regions showed elevated H3K9me3 levels relative to promoters and random regions, whereas this enrichment was less apparent in H1 and K562 cells (Fig. 3). These findings indicate that neocentromere landing regions generally reside within chromatin environments depleted of transcription-associated histone modifications and enriched for heterochromatic features, particularly H3K9me3. The observed cell-type variability further suggests that local chromatin context may contribute to neocentromere establishment and maintenance.

**Figure 3:**
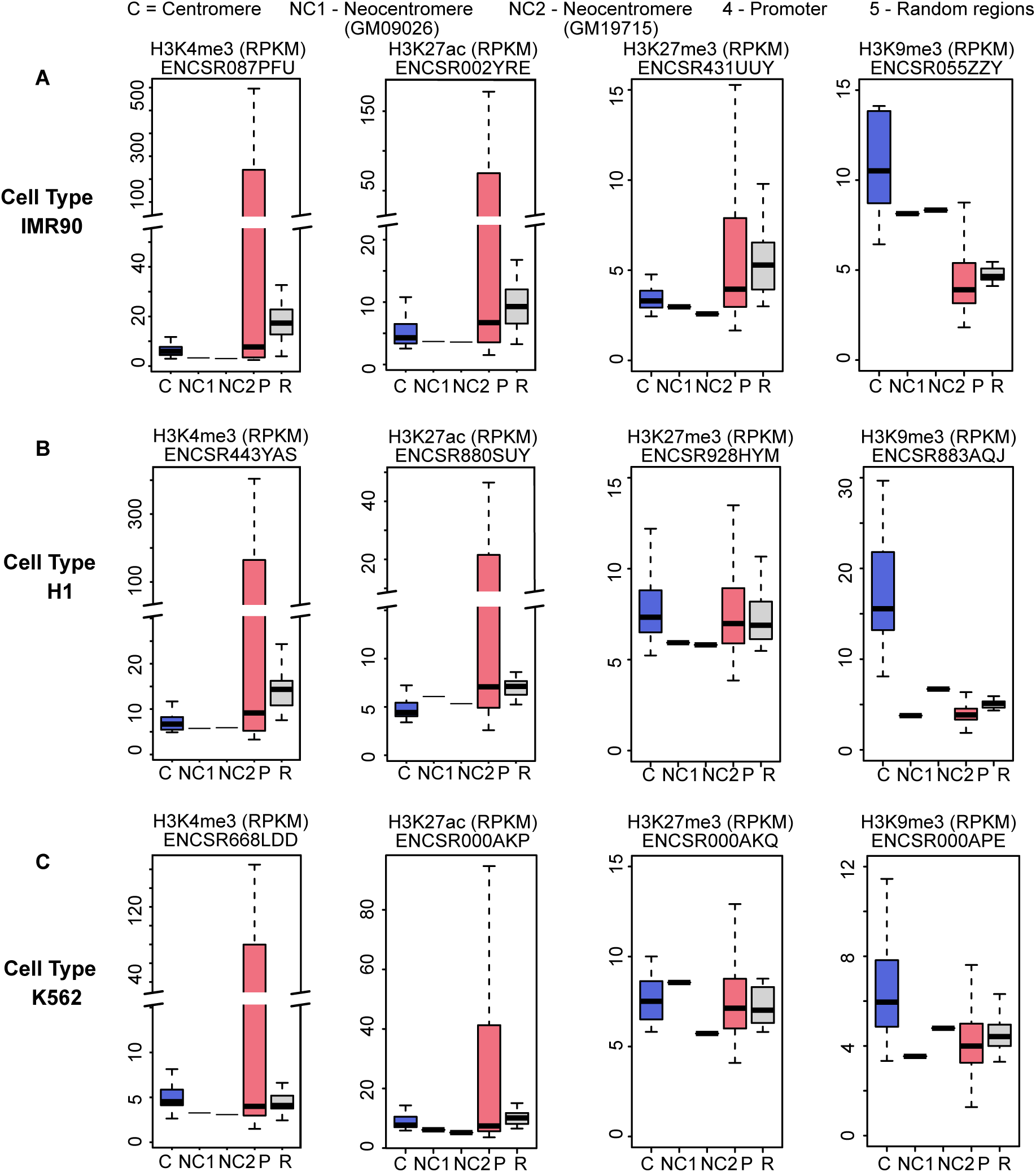
Depletion of active chromatin marks in neocentromere landing regions. **A –C**) Boxplots depict enrichment for H3K4me3, H3K27ac, H3K27me3, and H3K9me3 (RPKM) at centromeric regions, neocentromere landing region in human fibroblast cells (NC1 in GM09026), neocentromere landing region in lymphoblastoid cells (NC2 in GM19715), gene promoters, and random regions.

### Neocentromere landing regions are depleted of satellite DNA and reside within distinct epigenetic environments

Because canonical human centromeres possess large arrays of alpha-satellite DNA, we next examined whether similar repetitive sequence features are present at neocentromere landing regions. Analysis of repetitive and transposable element composition revealed marked differences between canonical centromeres and the two neocentromere regions examined (Fig. 4A). As expected, canonical centromeres were dominated by satellite-derived sequences, whereas both neocentromere landing regions lacked detectable satellite DNA. Instead, the neocentromeres were composed primarily of interspersed repetitive elements and unique genomic sequences. Notably, one neocentromere (NC2) exhibited substantial enrichment of LINE elements, whereas the second neocentromere (NC1) showed a repetitive element composition more similar to surrounding genomic background, indicating that specific classes of repetitive elements are not universally associated with neocentromere formation. To further characterize the local chromatin environment, we visualized histone modification profiles across both neocentromere regions (Fig. 4B). Consistent with the quantitative analyses shown in Figure 3, both loci were depleted of active promoter-associated marks, including H3K4me3 and H3K27ac, while exhibiting variable levels of H3K27me3 and H3K9me3. Together, these findings demonstrate that human neocentromeres can arise and persist in genomic regions devoid of alpha-satellite DNA and lacking a common repetitive sequence architecture. Instead, neocentromere formation appears to occur within transcriptionally inert chromatin environments, suggesting that local epigenetic states rather than specific repetitive DNA sequences influence whether genomic locations are more– or less-permissive to centromere establishment. However, the observational nature of these comparisons precludes direct testing of how chromatin features influence centromere formation, maintenance, and expansion.

**Figure 4:**
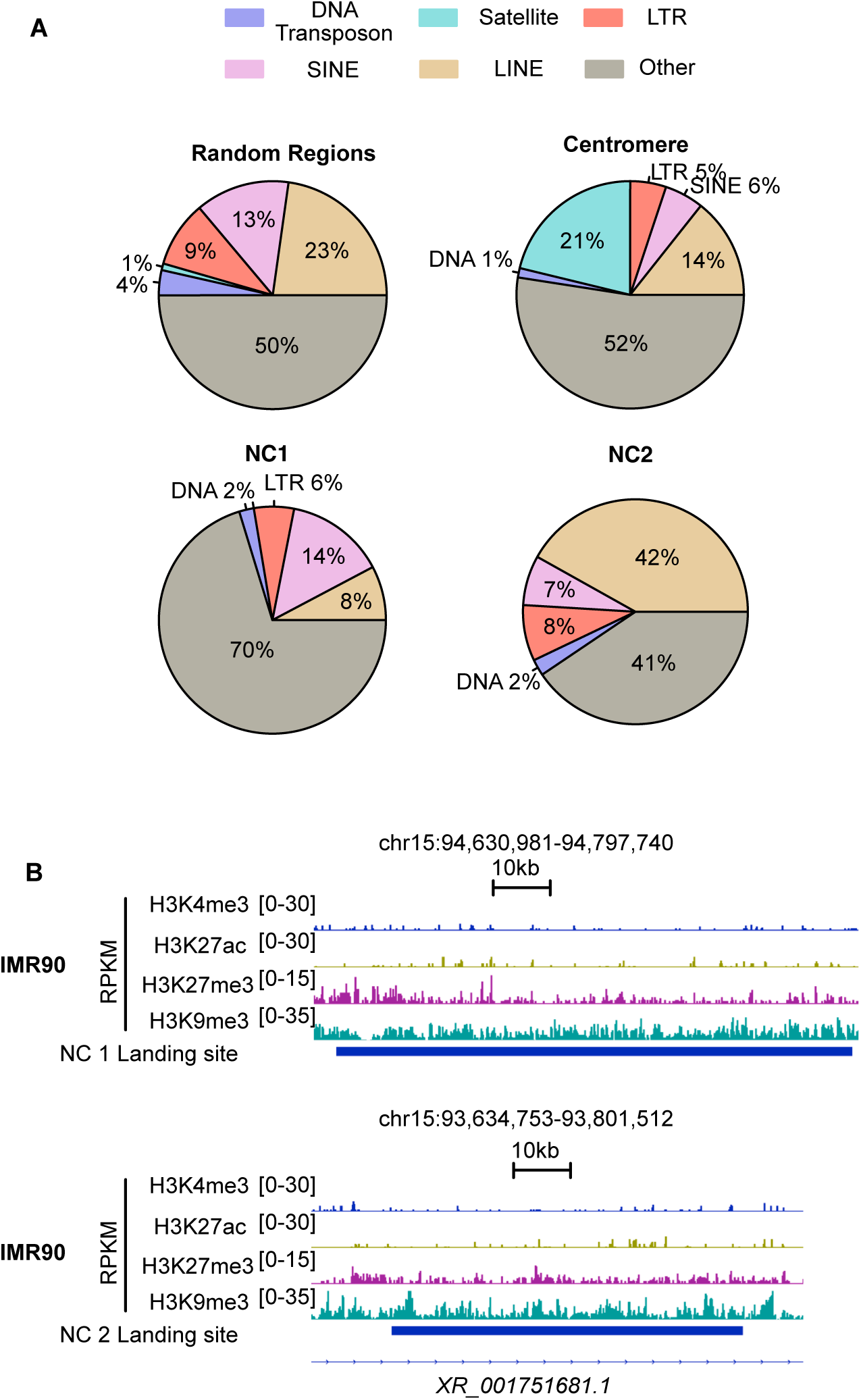
Distribution of genomic and epigenomic features in neocentromere landing regions. **A)** Pie chart showing the different classes of transposable elements distribution over random regions, centromeric regions, neocentromere landing region in human fibroblast cells (NC1), and neocentromere landing region in lymphoblastoid cells (NC2). **B)** Genomic browser track showing H3K4me3, H3K27ac, H3K27me3, and H3K9me3 enrichment (RPKM) over neocentromere landing region in human fibroblast cells (NC1), and neocentromere landing region in lymphoblastoid cells (NC2).

### Development of a genetically tractable monoallelic neocentromere system

We sought to develop a genetically tractable system in which the centromeric CENP-A domain could be selectively manipulated while preserving its native chromosomal context for mechanistic studies. Such direct genetic manipulation of centromeric chromatin in human cells is not possible at repetitive canonical centromeres. To overcome this limitation, we engineered a cell system in which the neocentromeric locus at 15q26.2, which is identical to endogenous sequences normally occurring on chromosome 15, is present as a single copy (only within the neocentromere) and can therefore be directly modified through genetic strategies. Because the primary fibroblast line GM09026 exhibited limited proliferative capacity and low transfection efficiency, we first generated a hybrid cell line, F026, through fusion of GM09026 cells with the human fibrosarcoma line HT1080 (Fig. 5A). Cytogenetic analyses demonstrated that the resulting hybrid cells retained the inv dup(15q) marker chromosome together with multiple normal chromosome 15 homologs. We then employed iterative rounds of CRISPR-Cas9-mediated chromosome engineering to remove all the non-centromeric alleles at 15q26.2 while preserving the neocentromeric CENP-A epi-allele (CA_epi) on the inv dup(15q) chromosome. This strategy generated the derivative line F026_65135, which contains a single remaining copy of the 15q26.2 region corresponding to the CA_epi allele (Fig. 5A). Interphase and metaphase FISH analyses confirmed the progressive reduction in copy number from G026 to F026 and ultimately to a monoallelic configuration at 15q26.2 in F026_65135 (Fig. 5B). Chromosome microarray analysis independently verified loss of the additional 15q26.2 alleles while retaining the neocentromeric locus (Fig. 5C). Furthermore, optical genome mapping confirmed the engineered 180-kb deletion on the inv dup(15q) chromosome and excluded unintended structural rearrangements at the targeted locus (Fig. 5D). Together, these studies established F026_65135 as a unique human cell system harboring a single genetically accessible neocentromere, enabling direct perturbation of centromeric chromatin and its surrounding genomic environment.

**Figure 5:**
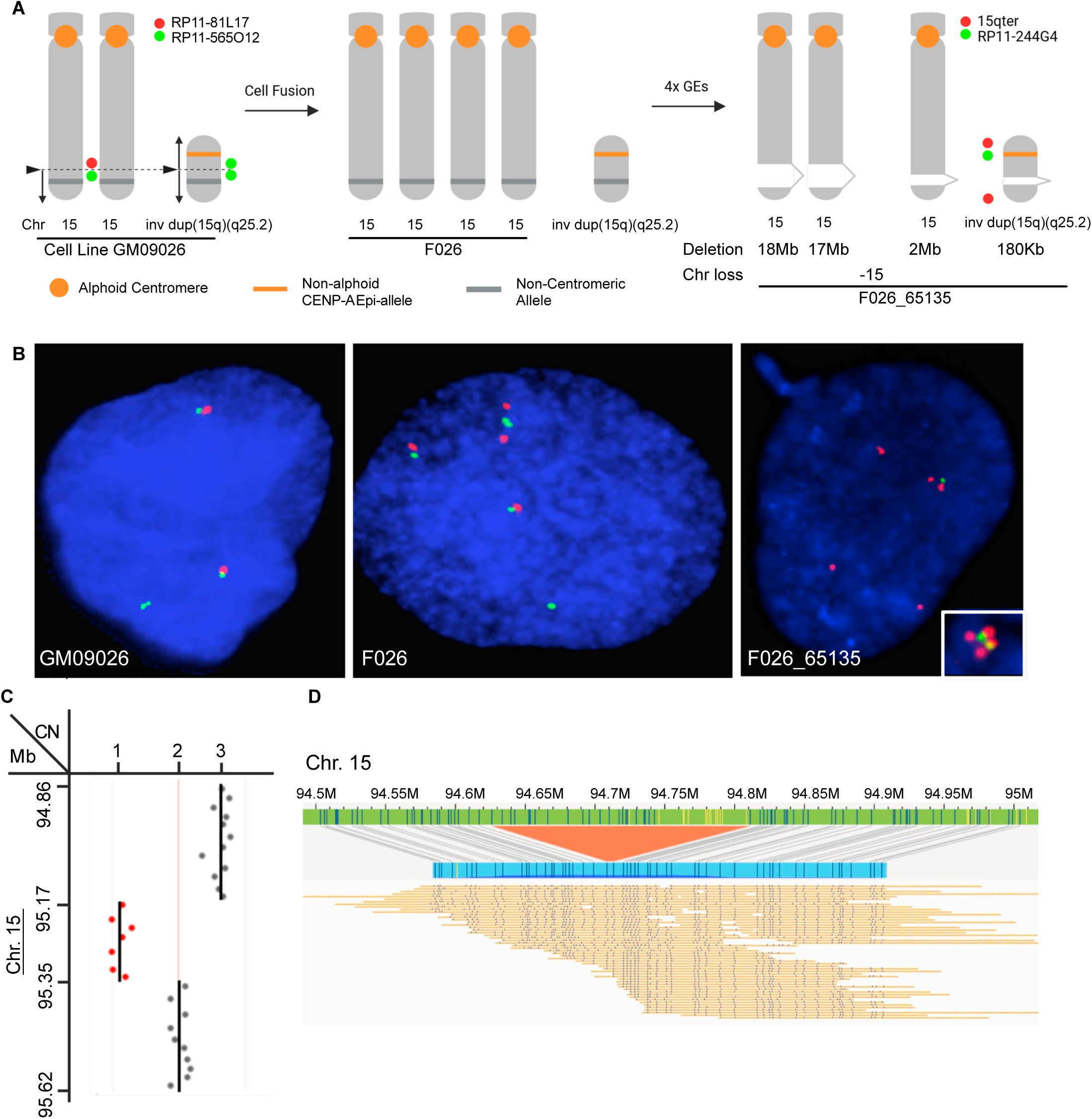
Generation of cell lines harboring a mono-allelic 15q26.2 region for direct genetic and epigenetic perturbation of the CENP-A epi-allele. **A)** Diagram of cell engineering and genome editing. G026 was fused with HT1080 to generate a line F026. Four rounds of genome editing were performed on F026 to produce the F026_65135 line that harbors 3 copies of chromosome 15 with 2 Mb, 17 Mb, and 18 Mb deletions surrounding 15q26.2, and one inv dup(15q) with 180 Kb deletion at 15q26.2, therefore leaving a monoallelic CENP-A epi-allele at 15q26.2. **B)** Interphase FISH confirmed copy numbers of chromosome 15 and 15q26.2 in cell line GM09026, F026_6, and F026_65135. Inset, metaphase FISH analysis of F026_65135. **C)** Chromosome microarray analysis (Agilent) shows copy number status around the CENP-A epi-allele at 15q26.2. **D)** Optical genome mapping analysis confirms the 180 kb deletion generated on the inv dup(15q).

### The monoallelic CENP-A epi-allele resides within a heterochromatic domain and is stably maintained through mitotic divisions

The monoallelic F026_65135 cell line provided an opportunity to examine the epigenetic landscape of the CENP-A epi-allele at allele-specific resolution, in contrast to the comparative analyses of neocentromere landing sites presented in Figures 3 and 4. We profiled chromatin accessibility and histone modifications across the 15q26.2 region in F026_65135 cells.

Integration of CUT&RUN and chromatin accessibility data revealed that the CENP-A epi-allele is embedded within a large (∼3 Mb) heterochromatic domain characterized by lack of ATAC-seq signals, enrichment of H3K9me3, and a paucity of active chromatin marks such as H3K4me3 (Fig. 6A). Despite residing within this transcriptionally repressive environment, the CENP-A chromatin domain remained sharply localized, suggesting that centromeric chromatin can be maintained within a broader heterochromatic compartment while retaining a distinct epigenetic identity. To determine whether the genomic position of the neocentromere remains stable over time, we performed CENP-A CUT&RUN analysis on multiple single cell-derived clones isolated from F026_65135. The resulting profiles showed highly concordant CENP-A occupancy patterns, with minimal variation in the location or extent of the CENP-A chromatin domain among clones (Fig. 6B). We next evaluated the mitotic stability of the inv dup(15q) chromosome by introducing an EGFP marker under the control of the EF1a promoter onto the chromosome and monitoring its retention by flow cytometry. Following prolonged propagation in culture, the vast majority of cells retained EGFP expression, indicating stable inheritance of the marker chromosome through successive cell divisions (Fig. 6C). This is consistent with the non-mosaic state of the marker chromosome observed in the parental lines (GM09026 and F026). Together, these findings demonstrate that the non-alphoid neocentromere is embedded within a stable heterochromatic domain and is faithfully maintained through mitosis, providing a robust platform for investigating the mechanisms governing centromeric chromatin maintenance and expansion.

**Figure 6:**
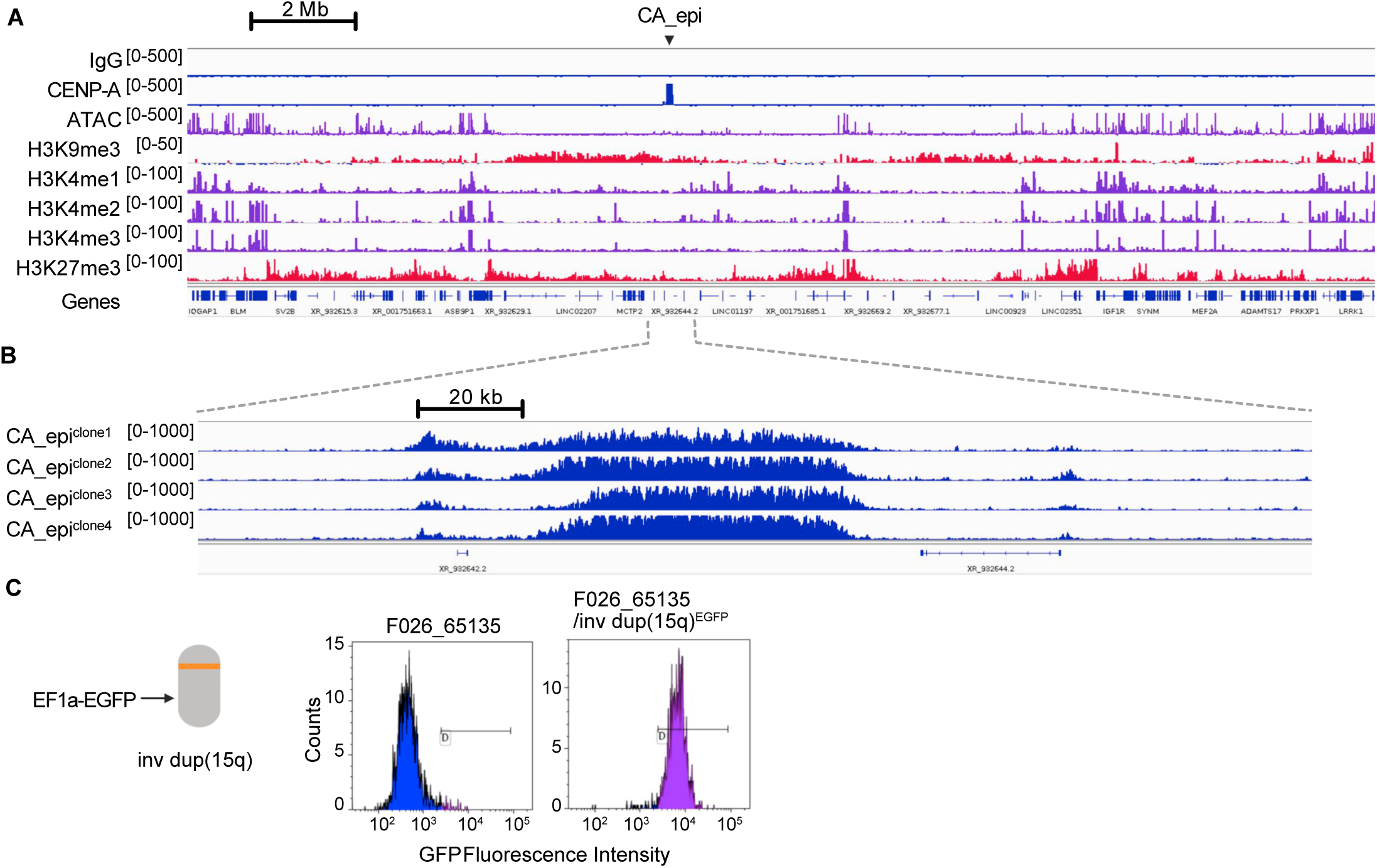
Epigenetic landscape surrounding the CA epi-allele and its stability in F026_65135. **A)** Profiling of epigenetic features in F026_65135 showed the CA epi-allele is situated within a heterochromatic domain (∼3 Mb) with closed chromatin (no ATAC signal), enriched in H3K9me3, and lack of H3K4me3. **B)** CUT&RUN analysis of CENP-A nucleosomes on single cell-derived clones from F026_65135 showed that the genomic location of the neocentromeric chromatin domain on the inv dup(15q) is stable. **C)** Flow cytometry analysis of F026_65135 and F026_65136/inv dup(15q)^EGFP^ showed the neocentromere is functional through mitotic cell divisions as no significant loss of the marker chromosome is observed.

### Truncation induces seed-driven expansion of centromeric chromatin and reveals mechanisms of domain size control

To better understand maintenance and establishment of functional centromeres, we engineered targeted deletions within the CENP-A epi-allele (CA_epi) and monitored subsequent changes in CENP-A occupancy. The unique nature of this 15q epi-allele enabled precise genetic manipulation. CRISPR-Cas9-mediated deletion of either 40 kb or 50 kb of the central CENP-A domain generated truncated centromeric chromatin seeds that remained associated with the inv dup(15q) chromosome (Fig. 7A). CUT&RUN analysis revealed that, in both CA_epi^del(40kb)#1^ and CA_epi^del(50kb)^ single-cell derived clones, the residual CENP-A chromatin expanded into adjacent naïve sequences that previously lacked detectable CENP-A occupancy. Expansion occurred beyond the deletion boundaries and restored a centromeric domain approaching the size of the original neocentromere, indicating that pre-existing CENP-A chromatin can serve as a seed for sequence-independent propagation of centromeric identity. In contrast, one independently derived CA_epi^del(50kb)^ clone failed to undergo chromatin expansion and subsequently lost the inv dup(15q) chromosome during propagation (CA_epi^loss^), demonstrating that successful restoration of centromeric chromatin is required for chromosome stability (Fig. 7A). To establish precise temporal control over this process, we developed a Cre-loxP system in which a 40-kb segment of the CA_epi domain could be excised in a regulatable manner. Cre-mediated recombination efficiently generated the expected deletion and similarly triggered expansion of the remaining CENP-A chromatin into newly juxtaposed naïve DNA (Fig. 7B), confirming that chromatin expansion is a reproducible response to centromeric chromatin damage rather than a consequence of CRISPR-mediated repair.

**Figure 7:**
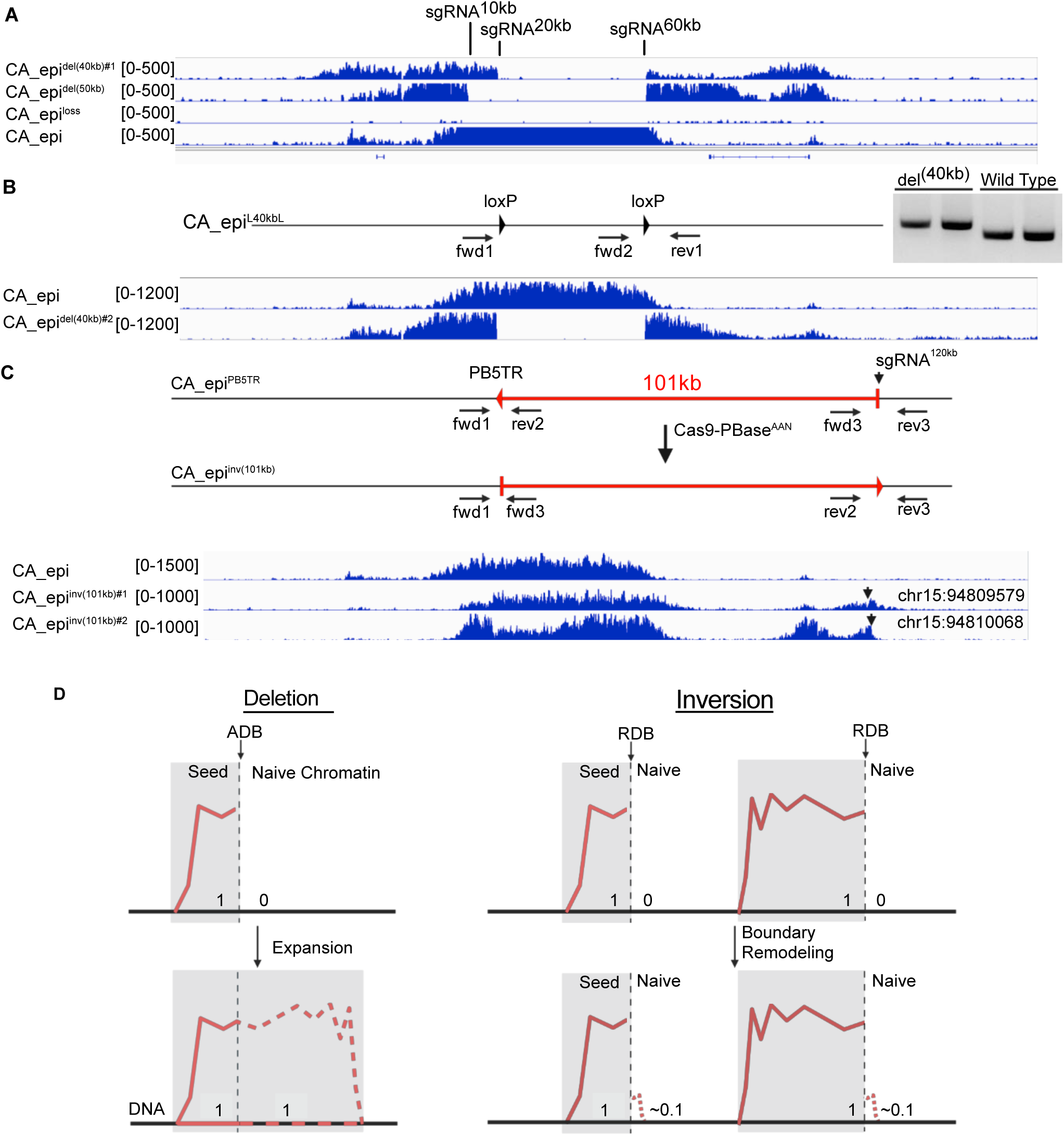
Epigenetic truncation and expansion of the CA epi-allele and chromosome inversion-mediated chromatin remodeling at domain boundaries. **A)** CRISPR-Cas9-mediated truncation induces seed-driven sequence-independent local CENP-A chromatin expansion. CA_epi^del(40kb)^ indicates a single cell derived clone harboring an expanded epi-allele following a 40kb deletion between sgRNA^20kb^ and sgRNA^60kb^. CA_epi^del(50kb)^ indicates a single cell derived clone harboring an expanded epi-allele following a 50kb deletion between sgRNA^10kb^ and sgRNA^60kb^. CA_epi^loss^ indicates a single cell derived clone with loss of the CA epi-allele following a 50kb deletion induced between sgRNA^10kb^ and sgRNA^60kb^ and subsequent loss of the inv dup(15q) marker chromosome. **B)** Cre-loxP-mediated 40 kb truncation induces chromatin expansion to adjacent naïve chromatin. The CA_epi^L40kbL^ allele harbors one loxP site at sgRNA^20kb^ and the other at sgRNA^60kb^. Cre recombinase triggers recombination between the two loxP sites, and induces a 40 kb deletion in a temporal manner. PCR analysis indicates the 40kb deletion. CUT&RUN analysis of single-cell derived clones show CENP-A chromatin expansion. **C)** 101 kb chromosome inversions induced by Cas9-PBase^AAN^ split the CENP-A chromatin domain and create new CENP-A/H3K9me3 chromatin boundaries. No significant CENP-A chromatin domain expansion was observed; however, modest boundary-associated redistribution of CENP-A occupancy occurred near the newly generated CENP-A/H3K9me3 interfaces. **D)** Diagram of a genetic system (left) that creates Active Domain Boundary (ADB) for studying seed-driven sequence-independent expansion of human centromeric chromatin. Diagram of a genetic system (right) that creates new CENP-A/H3 chromatin boundaries with no expansion potential for studying size control mechanism of human centromeric chromatin domain.

We next examined whether formation of new CENP-A chromatin domain boundaries is sufficient to trigger expansion. Two 101-kb inversions generated by Cas9-PBase^AAN^ split the local chromatin environment and created novel CENP-A/H3K9me3 interfaces without reducing overall centromeric chromatin mass (Fig. 7C). Under these conditions, we observed no substantial domain expansion. Instead, CUT&RUN analysis revealed modest boundary-associated redistribution of CENP-A occupancy near the newly created boundaries, likely attributed to restricted local drift of CENP-A nucleosomes. These findings indicate that the formation of new chromatin boundaries alone is insufficient to drive robust regenerative chromatin expansion. Rather, loss of centromeric chromatin mass appears to be the primary trigger for seed-driven expansion and restoration of centromere size, as reflected by the extent of the CENP-A chromatin domain. Importantly, these results distinguish regenerative expansion, triggered by loss of centromeric chromatin mass, from local boundary remodeling induced by creation of new chromatin interfaces.

## DISCUSSION

Centromeres are essential epigenetic structures that ensure faithful chromosome segregation, yet the mechanisms governing their establishment, maintenance, and size control remain poorly understood. Progress in the field has been constrained by the highly repetitive nature of canonical human centromeres, which precludes direct genetic manipulation of the underlying DNA. In this study, we developed a genetically tractable human neocentromere system based on a naturally occurring non-alphoid marker chromosome and used it to directly perturb centromeric chromatin in its native chromosomal context. A central finding is that partial deletion of a neocentromeric CENP-A domain triggers robust expansion of the remaining chromatin into adjacent naïve sequences. In multiple independently derived clones, residual CENP-A chromatin expanded beyond the truncation boundaries and re-established a domain approaching the size of the original neocentromere, whereas failure of expansion was associated with subsequent loss of the marker chromosome. These findings demonstrate that human centromeric chromatin possesses an intrinsic capacity to regenerate from residual CENP-A-containing chromatin seeds through sequence-independent chromatin expansion and provide experimental support for a two-step model of centromere formation in which acquisition of a CENP-A seed is followed by expansion into a mature centromeric domain capable of assembling a functional kinetochore. More broadly, the results suggest that centromeric chromatin is not maintained solely through the conservative template-guided replenishment mechanism that operates during normal cell cycles, but can also engage a regenerative mode when chromatin mass is reduced below a functional threshold. Such a mechanism may be important during spontaneous centromere damage, chromosome rearrangements, neocentromere formation, and centromere repositioning events observed in evolution and human disease. How such expansion/restoration occurs remains unknown.

Supernumerary marker chromosomes (SMCs) are structurally abnormal chromosomes associated with both constitutional disorders and cancer^14,15,18,36,37^. Most well-characterized human SMCs retain canonical alpha-satellite centromeres and therefore require little or no alteration of centromere identity for stable inheritance. In contrast, the constitutional inv dup(15q) chromosomes and the somatic inv dup(11q) chromosome described here belong to the much rarer class of non-alphoid SMCs that must acquire neocentromeres to ensure chromosome segregation^18^. Because these chromosomes arise naturally and are stably maintained in human cells, they provide a unique opportunity to investigate how centromeres are established, stabilized, and inherited in their native genomic context. All three inv dup(15q) chromosomes described in this study have breakpoints proximal to three different segmental duplication clusters at 15q24.1-3 (Fig. 1A), 15q25.2 (GM09026), 15q25.3 (GM19715), respectively. The relatively simple structure of these inv dup(15q) chromosomes, generated through a single Non-Allelic Homologous Recombination (NAHR) event, supports the hypothesis that neocentromere formation can occur on otherwise ordinary non-alphoid genomic DNA rather than requiring complex genomic rearrangements or cryptic insertions of alpha satellites^15^. These observations support the view that neocentromere establishment is fundamentally an epigenetic process and highlight naturally occurring non-alphoid SMCs as valuable models for investigating how centromeres are initiated, stabilized, and inherited in human cells. The rarity of non-alphoid SMCs in humans may reflect the low probability of acquiring a sufficiently large and stable CENP-A chromatin seed capable not only of establishing centromere identity but also of supporting subsequent chromatin expansion and maturation into a functional kinetochore.

Together, these results from both truncation and inversion experiments support a two-step model in which pre-existing CENP-A seeds direct sequence-independent chromatin expansion, while distinct boundary state switching mechanisms constrain domain growth and maintain centromere size homeostasis (Fig. 7D). Our results further provide insights into the longstanding question of how centromere size is regulated. Human centromeres contain relatively constant amounts of CENP-A despite substantial variation in the underlying DNA sequence content and repeat organization ^29,30,32^. Accordingly, centromere size is best considered in epigenetic terms, defined by the extent of the CENP-A chromatin domain rather than the amount of underlying centromeric DNA. Previous studies have suggested that centromere location and size are epigenetically inherited and subject to homeostatic regulation ^22,29,32^. The expansion observed following truncation is consistent with the existence of a size-control mechanism that restores a functional centromeric mass rather than simply maintaining existing nucleosome positions. In contrast, creation of new CENP-A/H3K9me3 boundaries by chromosomal inversion induced only modest boundary remodeling and local redistribution of CENP-A occupancy without triggering regenerative expansion. Together, these observations suggest that centromeres can exist in at least two functional states: a maintenance state bounded by restrictive domain boundaries that preserve a stable domain size, and a regenerative state characterized by Active Domain Boundaries (ADBs) that promote directional deposition of new CENP-A nucleosomes until centromeric chromatin mass is restored.

According to our model, these separate states underlie functional centromere formation, and the existence of alpha satellite DNA sequences would be a mere consequence of centromere expansion, rather than representing a driver – instead these repetitive sequences accumulate over time and expand in a non-deleterious manner.

Most importantly, our results indicate that the transition between these states is triggered by loss of centromeric chromatin mass rather than by the mere creation of new chromatin interfaces. Such a model provides a conceptual framework for understanding not only centromere maintenance but also neocentromere formation, centromere repositioning, and recovery from centromeric chromatin damage. Crucially, this regenerative mechanism may also provide a molecular explanation for classical genetic phenomena such as Position-Effect Variegation (PEV)^38^. In seminal Drosophila studies, X-ray irradiation-induced chromosomal inversions that placed euchromatic genes near pericentromeric heterochromatin resulted in stochastic gene silencing. We speculate that, if these radiation-induced breaks inadvertently severed portions of the centromere, the resulting loss of chromatin mass would trigger the active regenerative state. Under this model, sequence-independent expansion of the remaining centromeric seed — and the concomitant spreading of surrounding heterochromatic marks like H3K9me3 — is what drives the silencing of newly juxtaposed euchromatic genes. What was historically viewed as a passive “spreading” of heterochromatin may instead reflect an active, homeostatic program of centromeric chromatin restoration.

A major unanswered question raised by this work is how centromeres sense reductions in chromatin mass and activate the regenerative state. One possibility is that centromeric chromatin is monitored through quantitative feedback involving CENP-A nucleosomes and their associated CCAN components, such that loss of CENP-A density below a critical threshold triggers enhanced recruitment of factors required for CENP-A deposition and domain expansion, including components of the HJURP/MIS18 pathway. Alternatively, alterations in local chromatin architecture, heterochromatin organization, transcriptional activity, or higher-order chromatin interactions may serve as signals that promote regenerative expansion. More broadly, the molecular mechanisms that distinguish Active Domain Boundaries from Restrictive Boundaries remain unknown. Defining how centromeres detect chromatin loss, switch boundary states, and terminate expansion once an appropriate domain size has been restored will be essential for understanding how centromere size homeostasis is achieved while preserving long-term epigenetic inheritance.

An important advantage of the system described here is that it provides a direct experimental framework for identifying the factors that sense centromeric chromatin loss and activate the regenerative state. Centromeric chromatin expansion is a dynamic and probabilistic process: following truncation of the CENP-A domain, not all cells successfully restore a functional centromere. Cells that fail to expand the residual chromatin seed ultimately lose the GFP-marked inv dup(15q) chromosome, whereas cells that successfully regenerate centromeric chromatin retain the marker chromosome. This creates a powerful phenotypic selection system in which centromere regeneration can be quantitatively measured through chromosome retention. One could envision a similar evolutionary scenario in which a newly formed centromere seed arises on an acentric DNA fragment harboring essential or selectively advantageous genes. Analogous to retention of the GFP-marked chromosome in our experimental system, positive selection for these genes could favor stabilization of the chromosome through regenerative expansion of the centromeric chromatin, ultimately, contributing to *de novo* chromosome formation.

Moervoer, this platform is ideally suited for functional genetic screening. Unlike conventional centromere assays that monitor the static maintenance of pre-existing centromeres, this system converts regenerative centromere formation into a selectable phenotype with high dynamic range. Genome-wide or focused CRISPR-Cas9 screens can therefore be used to identify genes required for sensing centromeric chromatin loss, activating Active Domain Boundaries, promoting seed-driven chromatin expansion, or restoring centromere size homeostasis. Such studies have the potential to uncover previously unrecognized regulators of centromere formation, chromatin boundary dynamics, chromosome stability, and cellular responses to centromeric chromatin damage.

In addition to the factors that directly sense centromeric chromatin mass and regulate boundary-state switching, the surrounding chromatin environment may also influence centromere maintenance and expansion. The CENP-A epi-allele resides within a large H3K9me3-enriched domain that lacks active chromatin marks, resembling key features of canonical pericentromeric heterochromatin^18,39^. Similar enrichment of H3K9me3 was observed at independent neocentromere landing sites, although the degree of enrichment varied among cell types. These observations support a model in which heterochromatin provides a permissive environment for centromere stability and regeneration without directly specifying centromere identity. The absence of alpha-satellite DNA and the diversity of repetitive element composition among neocentromere regions further reinforce the concept that centromere identity is fundamentally epigenetic rather than sequence encoded. Future studies combining genetic perturbation with temporal epigenomic profiling during regeneration will enable direct testing of how heterochromatin, transcriptional activity, chromatin accessibility, DNA methylation, and three-dimensional genome organization contribute to centromeric chromatin expansion, domain boundary formation, and transitions between maintenance and regenerative states.

Beyond its utility for mechanistic studies, this system has broader implications for cancer biology and synthetic genomics. Chromosomal instability is a hallmark of many cancers and is frequently associated with aberrant expression of CENP-A, HJURP, and other kinetochore components^40–42^. The ability to experimentally induce centromeric chromatin damage and monitor subsequent repair or failure provides a unique opportunity to study cellular responses to compromised centromere function and to identify vulnerabilities associated with centromere maintenance. Likewise, understanding how small CENP-A seeds expand into mature centromeres may inform future strategies for constructing synthetic chromosomes. Current approaches to mammalian artificial chromosome generation remain limited by low efficiency and dependence on large arrays of alpha-satellite DNA. The demonstration that non-repetitive chromatin seeds can direct sequence-independent centromere expansion raises the possibility that efficient synthetic centromere construction may ultimately be achieved through epigenetic engineering rather than repetitive DNA assembly. By depositing a minimal, synthetically derived CENP-A seed onto a non-alphoid DNA backbone, researchers could exploit the host cell’s intrinsic centromere regenerative capacity to expand and mature a functional CENP-A domain *in situ*. This would dramatically streamline the synthesis of stable, heritable artificial chromosomes for durable gene replacement therapies^43^.

In summary, we describe a human genetic system that enables direct manipulation of centromeric chromatin and reveals a previously inaccessible regenerative program that restores centromeric chromatin following perturbation and maintains centromere size homeostasis. Our findings support a model in which centromere formation proceeds through two mechanistically distinct steps—seed acquisition followed by chromatin expansion—and suggest that Active Domain Boundaries (ADBs) drive restoration of centromeric chromatin until a characteristic domain size is achieved. By transforming centromere regeneration into a genetically selectable phenotype, this platform establishes a powerful framework for uncovering the molecular pathways that govern centromere inheritance, size control, chromatin boundary formation, and *de novo* centromere establishment in human cells.

## Supporting information

Reagents

## Acknowledgments

We thank Paula Vertino and Richard Burack for discussions, advice and critical comments on the manuscript; Ching-Hua Shih for providing bioinformatics support; and the members of the Cytogenetics, Array/OGM, and Molecular Diagnostic laboratories of URMC for providing quality genetic analysis. This work was supported by the Department of Pathology and Laboratory Medicine at University of Rochester Medical Center to B.Z., and was supported in part by Department of Defense grant W81XWH2210550 to A.P. and B.Z.. Additional support was provided to PJM from the National Institutes of Health grant R35GM137833.

## Author Contributions

BZ, ML, PJM, and SH conceived, designed, and performed the experiments. BZ, ML, PJM, and SH wrote the paper. All authors read and approved the final manuscript.

## Author Information

The authors declare no competing financial interests. Correspondence and requests for materials should be addressed to B.Z. lead contact and P.J.M.

## FIGURE LEGENDS

**Supplementary Figure 1:**
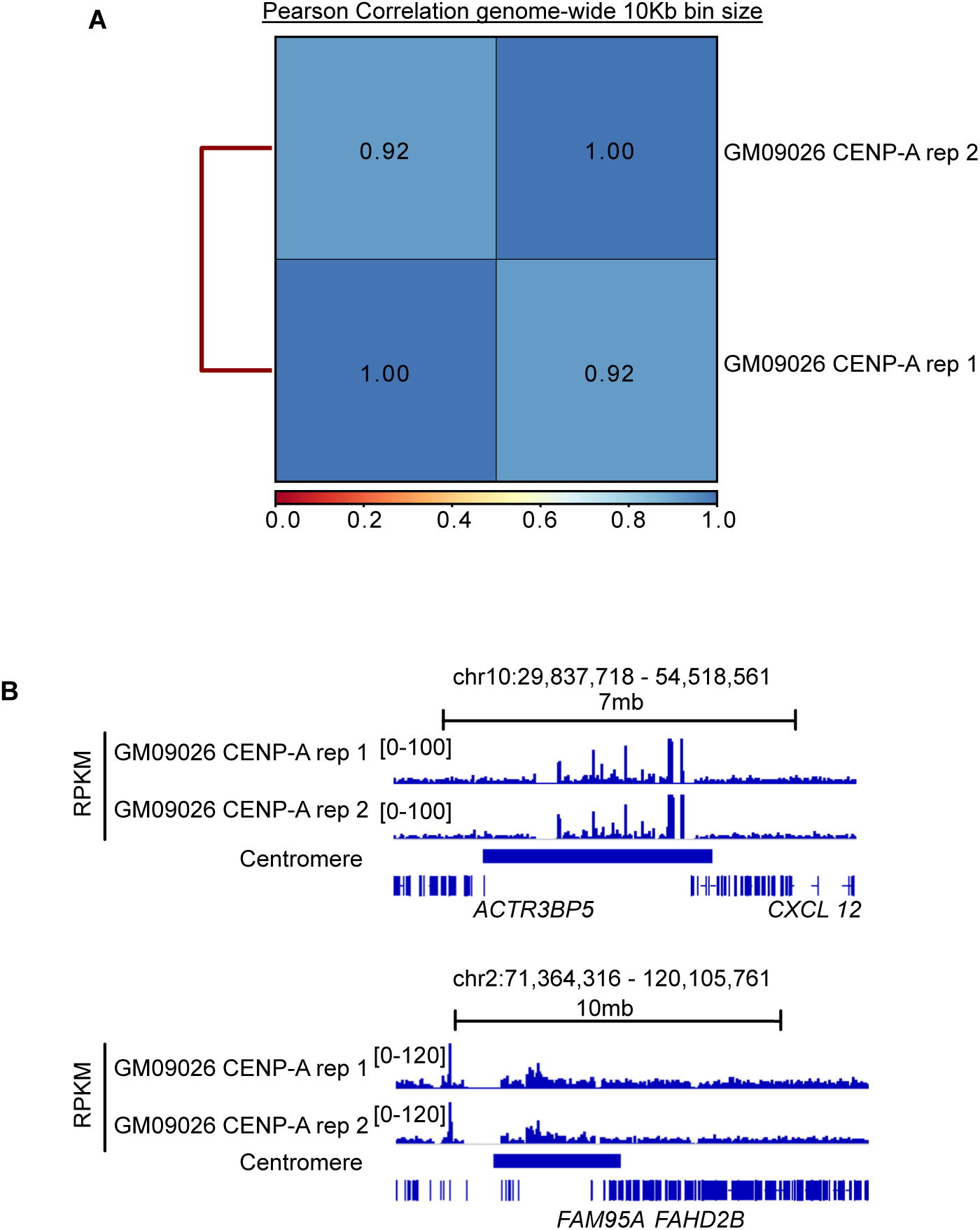
**A**) Pearson correlation heatmap showing the correlation between two replicates for CENP-A in GM09026 cell lines using 10kb bin-size. Pearson correlation coefficients are provided in the heatmap. **B)** Genomic browser track showing two replicates for CENP-A enrichment (RPKM) in GM09026 cell lines over centromeric regions in hg38 genome.

**Supplementary Figure 2:**
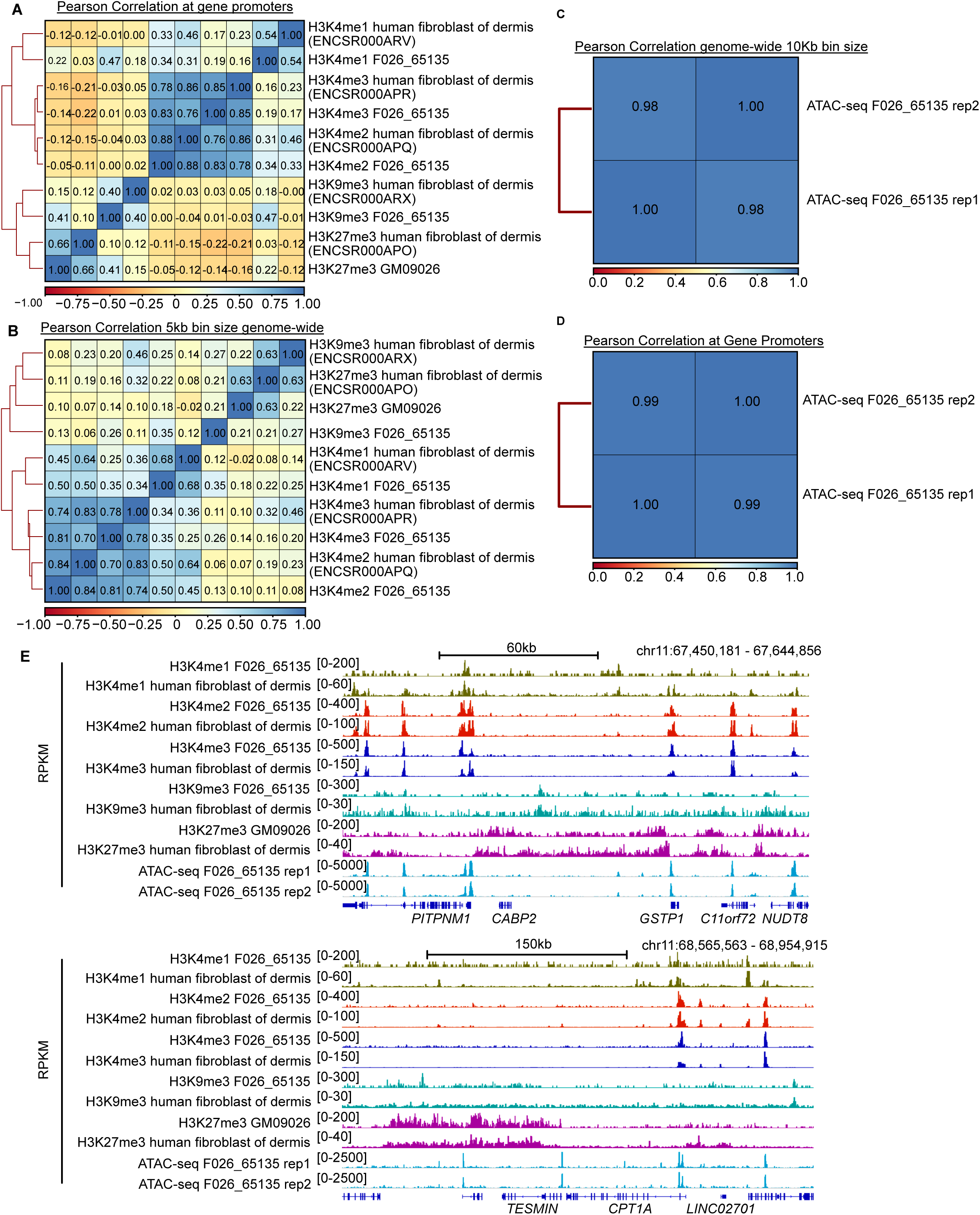
**A**) A matrix of Pearson correlation coefficients depicted in heatmap form, showing the correlation between ENCODE datasets and experimental data for H3K4me1, H3K4me2, H3K4me3, H3K9me3, and H3K27me3 in human fibroblast of the dermis, along with F026_65135 and GM09026. All genomic promoters were considered for assessing enrichment. **B)** The same type of Pearson correlation matrix provided in panel A, except the entire genome was broken into 5kb bins to assess enrichment. **C)** Pearson correlation heatmap showing the correlation between two replicates for ATAC-seq in F026_65135 using the entire genome broken into 10kb bins to assess enrichment. Pearson correlation coefficients are provided in the heatmap. **D)** The same type of Pearson correlation matrix provided in panel C, except genomic promoters were considered for assessing enrichment. **E)** Genomic browser track showing public available datasets and experimental data for H3K4me1, H3K4me2, H3K4me3, H3K9me3, and H3K27me3 enrichment (RPKM) in human fibroblast of dermis, F026_65135 and GM09026, respectively.

**Supplementary Figure 3:**
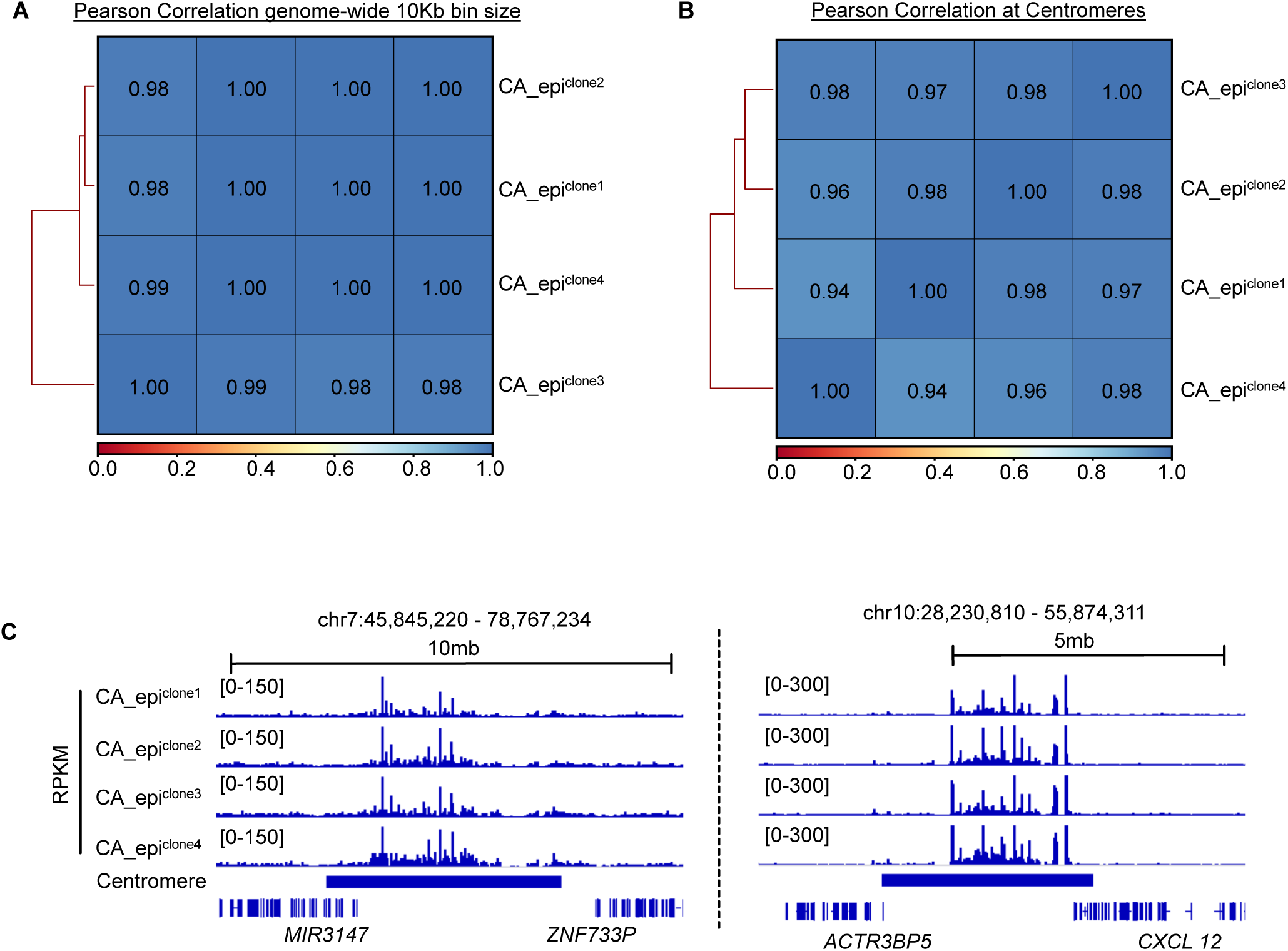
**A**) A heatmap showing Pearson correlation coefficients for the four replicates of CENP-A from single cell-derived clones, using the entire genome broken into 10kb bins to assess enrichment. are provided in the heatmap. **B)** The same type of Pearson correlation heatmap depicted in Panel A, except genome-wide centromeres were considered for enrichment. **C)** Genomic browser track showing four replicates of CENP-A enrichment focused on two true centromeres (RPKM) from single cell-derived clones.

**Supplementary Figure 4:**
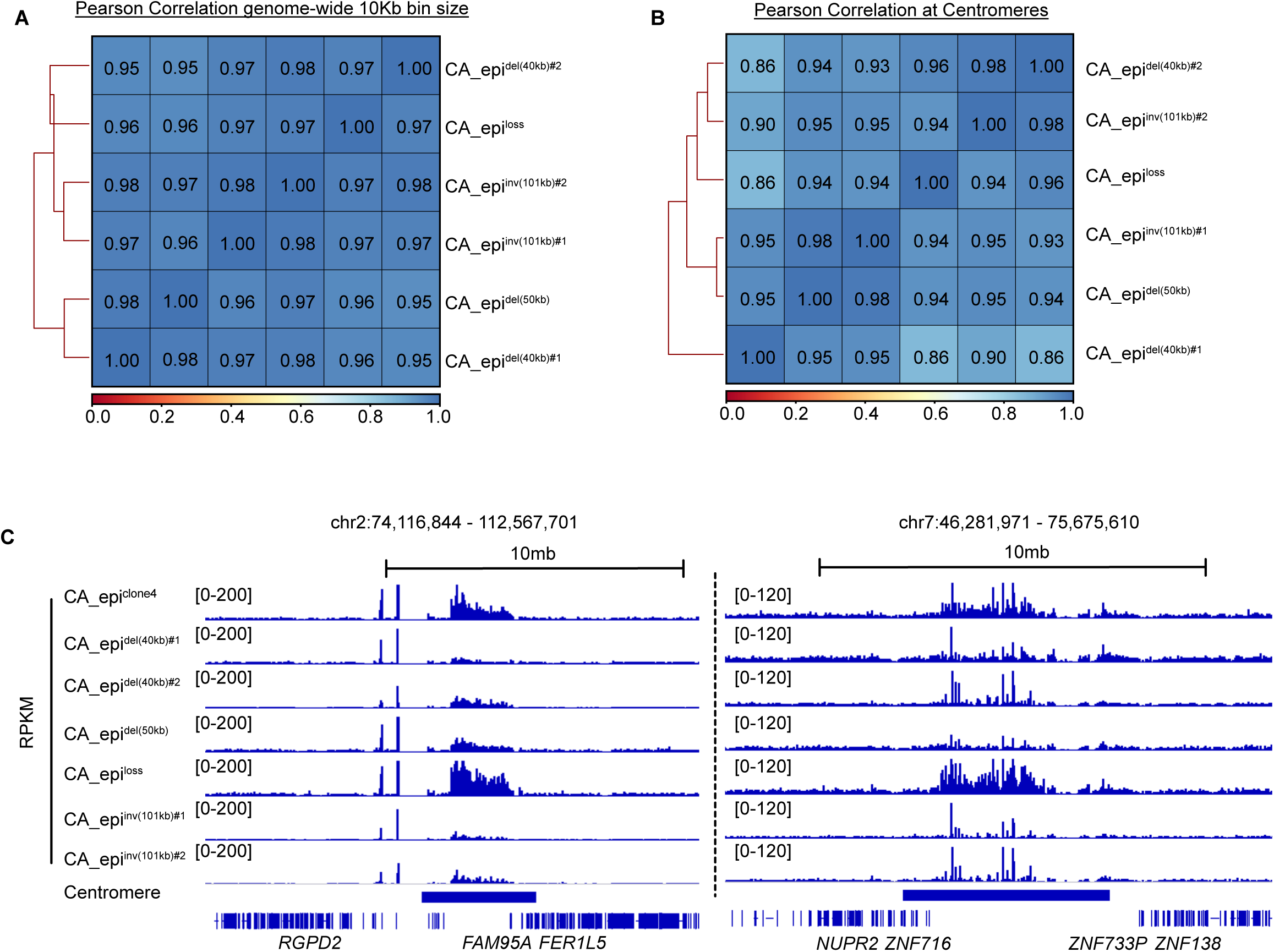
**A**) A heatmap showing Pearson correlation coefficients measurement of CENP-A enrichment across clonal cell lines in which distinct portions of the neocentromere were deleted. The entire genome broken into 10kb bins was used to assess enrichment. **B)** The same type of Pearson correlation heatmap depicted in Panel A, except genome-wide centromeres were considered for enrichment. **C)** Genomic browser track showing four replicates of CENP-A enrichment focused on two true centromeres (RPKM), using clonal cell lines in which distinct portions of the neocentromere were deleted.

## METHODS

### Copy Number Analysis using Chromosome MicroArray (CMA)

Genomic DNA was isolated from peripheral blood lymphocytes or cultured cells using QIAamp DNA Blood Mini Kit (Qiagen, CA). A Nanodrop ND-2000 spectrometer (ThermoFisher Scientific, MA) was used for determination of DNA concentrations. Chromosome microarray analysis was performed with an input amount of 500 ng of genomic DNA using either Applied Biosystems CytoScan HD arrays (ThermoFisher Scientific, MA) or Agilent SurePrint G3 Human CGH Microarray 4 × 180 K platform (Agilent Technologies, CA) following manufacturers’ protocols ^44,45^. For CytoScan HD arrays, data were analyzed using Genotyping Console v4.0 (Affymetrix, CA) with an *in silico* reference library, and visualized by Chromosome Analysis Suite (ChAS; ThermoFisher Scientific, MA). For Agilent arrays, commercially available pooled male DNA (Promega, WI) was used as control, and data were analyzed and visualized using the Agilent CytoGenomics 4.0 software (Agilent Technologies, CA). All genomic coordinates are based on the Human GRCh38/hg38 Genome Assembly. CMA data were used for characterization of marker chromosomes, confirmation of CRISPR-engineered deletions, and evaluation of genome-wide copy number integrity in engineered cell lines.

### Cytogenetic analysis

Conventional chromosome analysis was performed using standard cytogenetic methods^44^. Cells were cultured under appropriate conditions, arrested in metaphase with colcemid, exposed to hypotonic treatment, and fixed in methanol:acetic acid (3:1). Chromosomes were analyzed by G-banding using trypsin digestion and Wright’s staining (GTW). Karyotypes were interpreted and reported according to the International System for Human Cytogenomic Nomenclature (ISCN 2024). At least 20 metaphase cells were analyzed when available.

### Fluorescence In Situ Hybridization (FISH) and ImmunoFluorescence (IF) Analysis

Fluorescence in situ hybridization (FISH) was performed on metaphase chromosomes and interphase nuclei using commercially available centromeric and locus-specific probes or custom bacterial artificial chromosome (BAC)-derived probes (Abbott Molecular, IL; Empire Genomics, NY). Probe hybridization, post-hybridization washing, and signal detection were performed using standard protocols. Slides were counterstained with DAPI and analyzed using a fluorescence microscope equipped with a CytoVision imaging system (Leica Biosystems, Germany).

Combined immunofluorescence and fluorescence in situ hybridization (IF/FISH) was performed to simultaneously visualize centromeric chromatin and underlying genomic loci^46^. Briefly, cells were cultured with 100 ng/ml Colcemid (Invitrogen; Thermo Fisher Scientific, MA) for 2 hours, and metaphase cells were harvested by mitotic blow-off, washed twice with PBS, and treated with pre-warm hypotonic buffer (75 mM KCl) at 37 degree for 30 min, and then centrifuged and resuspended in hypotonic buffer. 2-4×10^4^ cells in 500 ul were concentrated onto each ethanol washed slide at 1,800 rpm for 10 min using a Shandon Cytospin 4 Cytocentrifuge (Thermo Fisher Scientific, MA). Cells were prefixed with 2% (v/v) formaldehyde at room temperature for 10 min. Chromosome spreads were first immunostained with anti-CENP-A antibody (1:100) followed by a secondary antibody (Goat anti-mouse, 1:300). Cells were then fixed in 4-6%(v/v) formaldehyde for 15 min, then denatured and hybridized with locus-specific BAC probes, and countered stained with DAPI. Fluorescence signals were captured and analyzed using the CytoVision imaging system (Leica Biosystems, Germany). Cytogenetic analyses, including G-banding, FISH, and IF/FISH, were used to characterize constitutional and somatic marker chromosomes, define chromosomal breakpoints, validate engineered genomic deletions, localize the CENP-A epi-allele, and assess chromosome stability in engineered cell lines. All the antibodies and FISH probes are provided in in the supplementary file (Sup. File 1: Reagents Information).

### Plasmid Construction and sgRNA Cloning

All the sgRNAs were cloned into the LRG2.1T (Addgene: #108098) vector with an optimized sgRNA scaffold backbone using a *BsmBI* restriction site^47^. All the other vectors, including the Cas9, Cre, Hygro, Puro, Neo, hyPBase expression vectors and the PITCh insert vectors, were cloned using Gibson Assembly (NEB #E2611). Mutations for hyPBase^AAN^ was introduced using Q5 site-directed mutagenesis (NEB #E0554). All the oligonucleotide sequences used for vector construction are provided in the supplementary file (Sup. File 1: Reagents Information).

### CRISPR-Cas9-based genome deletions

CRISPR-Cas9-based genome editing ^48,49^ was performed to generate targeted deletions at the 15q26.2 locus and to engineer derivative cell lines carrying a monoallelic CENP-A epi-allele. Single-guide RNAs (sgRNAs) were designed to target unique non-repetitive sequences flanking the regions of interest. For each deletion, pairs of sgRNAs were cloned into the sgRNA expression vector LRG2.1T. Cells were co-transfected with sgRNA vectors and LentiCas9-Blasticidin construct using Fugene HD (Promega, WI) with standard transfection conditions optimized for the F026 cell line. Transfected cells were enriched by blasticidin pulse selection for 48 h after splitted onto new dishes.

Single-cell-derived clones were isolated by ring-picking or limiting dilution, and expanded for downstream analysis. Targeted deletions were screened by junction PCR using primers flanking the predicted deletion breakpoints and confirmed by Sanger sequencing when applicable. Positive clones were further validated by fluorescence in situ hybridization, chromosome microarray analysis, and/or optical genome mapping to confirm the intended genomic deletion and exclude unintended large-scale structural rearrangements. This strategy was used for iterative removal of the non-centromeric alleles at 15q26.2, generation of the F026_65135 monoallelic CENP-A epi-allele cell line, and targeted truncation of the CENP-A chromatin domain for analysis of seed-driven centromeric chromatin expansion. All the oligonucleotide sequences used for PCR genotyping are provided in the supplementary file (Sup. File 1: Reagents Information).

### Precise Integration into Target Chromosome (PITCh)-mediated insertions

PITCh was performed to introduce donor constructs into the 15q26.2 locus^50^. Briefly, donor plasmids containing the desired insertion cassette flanked by short microhomology sequences were co-transfected with LentiCas9-Blasticidin-sgPITCh (modified from plasmid Addgene #52962) and LRG2.1T sgRNA expression constructs. Cas9-mediated double-strand breaks generated at the target locus were repaired through microhomology-mediated end joining (MMEJ), resulting in targeted integration of the donor cassette at the intended genomic site. Single-cell-derived clones were isolated and screened and confirmed as described above in the section of CRISPR-Cas9-based genome deletions. PITCh-mediated integration was used to introduce loxP sequences into the CENP-A epi-allele for temporal induction of CENP-A chromatin truncation, and PB5TR cassettes for subsequent FiCAT/Cas9-PBase^AAN^-mediated chromosomal inversion experiments. All the plasmid construction information and all the oligonucleotide sequences used for PCR genotyping are provided in the supplementary file (Sup. File 1: Reagents Information).

### FiCAT/Cas9-hyPBase^AAN^-mediated chromosomal inversion

Large chromosomal inversions were generated using a modified genome engineering method modified from FiCAT (Find and Cut-And-Transfer) as previously described^51,52^. Briefly, a DNA fragment, consisting of piggyBac transposon 5’ inverted terminal repeats (ITRs) (PB5TR) flanked by locus-specific micro-homology sequences, was introduced into the middle of the CENP-A epi-allele at 15q26.2 in F026_65135 using the PITCh approach. sgRNAs targeting sequences ∼100 kb downstream of the PB5TR insertion site were designed. Cells were co-transfected with LRG2.1T-sgRNA expression and Lenti_Cas9_PBase^AAN^_Blast constructs^52^.

PBase^AAN^ is an excision competent/integration defective (Exc+Int−) transposase with R372A/K375A/D450N mutations^52^. Simultaneous Cas9-mediated cleavage at the sgRNA target site and PBase^AAN^–mediated cleavage at the PB5TR site generated chromosomal inversions through reorientation and non-homologous end joining of the intervening genomic segment in between. Following transfection, positive single-cell-derived clones were isolated, expanded, and analyzed. All the plasmid construction information and all the oligonucleotide sequences used for PCR genotyping are provided in the supplementary file (Sup. File 1: Reagents Information).

### Optical genome mapping analysis

Optical genome mapping (OGM) was performed using the Bionano Saphyr system according to the manufacturer’s protocols^53^. Ultra-high molecular weight genomic DNA was isolated from 1 to 1.5×10^6^ cultured cells (Bionano, CA). A total of 750 ng of genomic DNA, was labeled enzymatically, conjugating fluorophores to the target 6-mer CTTAAG without cleaving the phosphate backbone. Long, labeled DNA strands were then counterstained with an intercalating dye, homogenized in buffer, and loaded into flowcells of Saphyr G2.3 chips (Bionano). A target throughput of 800 Gbp was achieved per flowcell. Completed data sets were then assessed for analytical quality control, targeting 160× effective coverage of GRCh38/hg38, with ≥70% of molecules ≥150 kbp aligning (map rate) at an N50 of ≥230 kbp (N50 is defined by the length of the shortest molecule for which equal and longer molecules account for 50% of the total data). Completed data sets were uploaded to a central server running Bionano Access version 1.7 or 1.7.1. (hereafter, v1.7.x) for analysis with Bionano Solve 3.7 or 3.7.1. (hereafter, v3.7.x). GRCh38/hg38 was selected as reference, and bioinformatic analyses were launched from Access to Solve pipeline software (Bionano, CA). Copy number variants and structural variants were called using the rare variant or *de novo* assembly pipeline. OGM was used to characterize marker chromosome structure, confirm engineered deletions at 15q26.2, validate chromosome rearrangements, and assess the absence of unexpected large-scale structural alterations in engineered cell lines and the absence of unexpected repeat insertion at 15q26.2 in GM09026.

### Flow cytometry

Flow cytometry was performed to monitor retention or loss of the EGFP-marked inv dup(15q) chromosome and to quantify chromosome stability. Cells were harvested at the indicated time points, washed with phosphate-buffered saline (PBS), and resuspended as single-cell suspensions in PBS containing fetal bovine serum. Cell aggregates and debris were excluded by forward– and side-scatter gating, and viable single cells were analyzed for EGFP fluorescence. Parental cells lacking EGFP were used to establish background fluorescence and define the EGFP-negative gate. For each sample, at least 10,000 single-cell events were collected when available. Data were analyzed using standard flow cytometry software. This assay was used to assess mitotic stability of the inv dup(15q) chromosome and to quantify chromosome loss following perturbation of the CENP-A epi-allele.

### CUT&RUN and ATAC-seq

CUT&RUN was performed as previously described using the EpiCypher CUTANA CUT&RUN protocol and reagents, with minor modifications (^35,54,55^ and epiCypher, NC). Briefly, cells were harvested, washed, and immobilized on concanavalin A–coated magnetic beads.

Permeabilized cells were incubated with primary antibodies against CENP-A, H3K4me1, H3K4me2, H3K4me3, H3K9me3, H3K27me3, or control IgG at 4 °C over night. After washing, cells were incubated with protein A/G–micrococcal nuclease fusion protein (SKU: 15-1016, EpiCypher, NC), followed by calcium-dependent activation to release antibody-targeted chromatin fragments. Reactions were stopped by addition of chelating buffer, and released DNA fragments were recovered from the supernatant and purified. Sequencing libraries were prepared from purified CUT&RUN DNA using NEBNext® UltraTM II Library Prep Kit (E7645S, New England Biolabs, MA) and subjected to paired-end sequencing (MiSeq, Illumina, CA). All the antibodies are provided in in the supplementary file (Sup. File 1: Reagents Information).

ATAC-seq library prep was performed as previously described^56^. Briefly, 50,000 cells were washed three times with cold PBS, collected by centrifugation then lysed in lysis buffer (10 mM Tris-HCl, pH 7.4, 10 mM NaCl, 3 mM MgCl2, 0.1% NP-40). After purification of nuclei, transposition was performed with Tn5 transposase from Nextera DNA Library Prep Kit (Illumina, catalog # FC-121-1030). Purified DNA was then ligated with adapters, amplified and size selected for sequencing. Libraries were sequenced with Illumina HiSeq3000 or Illumina NovaSeq6000.

Replicates were performed for each sample. Reads were aligned to the human reference genome Assembly GRCh38/hg38 using Bowtie2, and duplicates were removed using Picard. Read count normalization (RPKM) was performed on alignment files. Normalized read density was calculated to identify CENP-A-enriched domains and histone modification profiles and chromatin accessibility across the 15q26.2 region and the genome. Genome track visualized in Integrative Genomics Viewer (IGV). CUT&RUN was used to map the CENP-A epi-allele, assess its stability in single-cell-derived clones, and quantify CENP-A chromatin redistribution following genetic perturbation. ATAC-seq was used to assess the state of the CENP-A chromatin domain and its adjacent chromatin at 15q26.2.

### Processing of ChIP-seq datasets

The bam files for ChIP-seq data of H3K4me1, H3K4me2, H3K4me3, H3K27ac, H3K27me3, and H3K9me3 in IMR90, H1, K562 and human fibroblast of dermis cell lines were downloaded from the ENCODE project using the corresponding accession numbers: H3K4me3 in IMR90, ENCSR087PFU; H3K27ac in IMR90, ENCSR002YRE; H3K27me3 in IMR90, ENCSR431UUY; H3K9me3 in IMR90, ENCSR055ZZY; H3K4me3 in H1, ENCSR443YAS; H3K27ac in H1, ENCSR880SUY; H3K27me3 in H1, ENCSR928HYM; H3K9me3 in H1, ENCSR883AQJ; H3K4me3 in K562, ENCSR668LDD; H3K27ac in K562, ENCSR000AKP; H3K27me3 in K562, ENCSR000AKQ; H3K9me3 in K562, ENCSR000APE. H3K4me1 in human fibroblast of dermis: ENCSR000ARV; H3K4me2 in human fibroblast of dermis: ENCSR000APQ; H3K4me3 in human fibroblast of dermis: ENCSR000APR; H3K9me3 in human fibroblast of dermis: ENCSR000ARX; H3K27me3 in human fibroblast of dermis: ENCSR000APO. Picard (v2.12.0) BuildBamIndex was used to generate index files for each bam file. Deeptools (v3.5.1) bamCoverage was used to convert bam files to bigwig files with read counts normalization (RPKM) and 10bp bin size. Merged bigwig file were processed by averaging of merging two technical replicates using bigWigMerge (v2.10).

### Genomic Distribution Analysis

The centromere coordinates in human genome (hg38) were downloaded from the UCSC Table Browser and expanded by 1 Mb on each side to include pericentromeric regions. Random background regions were generated using bedtools (v2.30.0) shuffle. Bedtools intersect was used to calculate the number of overlapping base pairs between each transposable elements class with centromeric, neocentromeric and random regions. The overlapping base pairs were divided by total length of each region set to calculate percentage.

### CUT&RUN and ATAC-seq data quality assessment

Data quality was evaluated using multiple complementary approaches. For datasets generated with biological and technical replicates, reproducibility was assessed by calculating Pearson correlation coefficients using genome-wide binned read counts or signal intensity over centromeric regions, as appropriate. For histone modification profiling, CUT&RUN datasets were further benchmarked against publicly available ChIP-seq datasets from human dermal fibroblasts by comparing genome-wide signal distributions and enrichment patterns over gene promoters. Representative genome browser tracks were also inspected to confirm concordance between biological replicates and with published reference datasets. Comprehensive quality assessment of all CUT&RUN and ATAC-seq datasets is provided in Supplementary Figures 1– 4.

### Plotting and visualization

All the Pearson Correlation plots were generated using Deeptools multibigwigSummary and plotCorrelation (v3.5.1). R studio with defulat settings was used to generate pie chart and boxplots. The Intergrative Genomics Viewer (IGV v2.16.0) was used to visulize the genomic tracks. Texts, font sizes and colors were adjusted in Affinity Designer.

